# Neural measures of subsequent memory reflect endogenous variability in cognitive function

**DOI:** 10.1101/576173

**Authors:** Christoph T. Weidemann, Michael J. Kahana

## Abstract

Human cognition exhibits a striking degree of variability: Sometimes we rapidly forge new associations whereas at other times new information simply does not stick. Correlations between neural activity during encoding and subsequent retrieval performance have implicated such “subsequent memory effects” (SMEs) as important for understanding the neural basis of memory formation. Uncontrolled variability in external factors that also predict memory performance, however, confounds the interpretation of these effects. By controlling for a comprehensive set of external variables, we investigated the extent to which neural correlates of successful memory encoding reflect variability in endogenous brain states. We show that external variables that reliably predict memory performance have relatively small effects on electroencephalographic (EEG) correlates of successful memory encoding. Instead, the brain activity that is diagnostic of successful encoding primarily reflects fluctuations in endogenous neural activity. These findings link neural activity during learning to endogenous states that drive variability in human cognition.

---

The capacity to learn new information can vary considerably from moment to moment. We all recognize this variability in the frustration and embarrassment that accompanies associated memory lapses. Researchers investigate the neural basis of this variability by analyzing brain activity during the encoding phase of a memory experiment as a function of each item’s subsequent retrieval success. Across hundreds of such studies, the resulting contrasts, termed subsequent memory effects (SMEs), have revealed reliable biomarkers of successful memory encoding (Paller & Wagner, 2002; Kim, 2011; Hanslmayr & Staudigl, 2014).

A key question, however, is whether the observed SMEs indicate endogenously varying brain states, or whether they instead reflect variation in external stimulus- and task-related variables, such as item difficulty or proactive interference, known to strongly predict subsequent memory (Kahana, Aggarwal, & Phan, 2018). It is tempting to attribute SMEs to endogenous factors affecting encoding processes and/or to specific experimental manipulations (such as encoding instructions) aimed at directly affecting these processes (Fellner, Bäuml, & Hanslmayr, 2013; Hanslmayr, Spitzer, & Bäuml, 2009; Hanslmayr & Staudigl, 2014). At the same time, some of the strongest predictors of recall performance are characteristics of individual items (e.g., pre-experimental familiarity or position in the study list; DeLosh & McDaniel, 1996; Merritt, DeLosh, & McDaniel, 2006; Murdock, 1962) whose effects are difficult to distinguish from those of endogenous factors, given that the successful retrieval of individual items is not under direct experimental control. Such idiosyncratic effects are therefore serious confounds in SME analyses and the relative contributions of endogenous and external factors to the SME have yet to be established.

Here we approach these challenges in two ways using a large free-recall data set comprising 97 individuals who each had their EEG recorded while they studied and recalled 24 word lists in each of at least 20 experimental sessions that took place over the course of several weeks. As shown in Figure 1a, the presentation of each list was followed by a distractor task and a free recall test. Each list contained 24 words and the same 576 words (24 words in 24 lists) were presented in each session, but their assignment to lists, and serial positions within lists, varied (we also refer to individual word presentations as “items” irrespective of the word identities). Our first approach closely builds on standard SME analyses that compute a contrast for neural activity during each item’s presentation in the study list. Rather than only predicting subsequent memory as a binary variable, however, we also statistically accounted for a comprehensive set of external factors that correlate with recall performance and computed SMEs for the corresponding residuals. Whereas residuals of statistical models are often treated as “noise,” here they reflect variability in recall performance that was not accounted for by external factors and they thus highlight contributions of endogenous factors to the SME. Comparing SMEs for these residuals with the standard item-level SME predicting binary retrieval success thus allowed us to estimate the relative contributions of endogenous neural variability and external factors to the SME (to the extent that SMEs are driven by external factors, SMEs should be absent when the effects of these external factors are statistically removed from recall performance).

**Figure 1.**
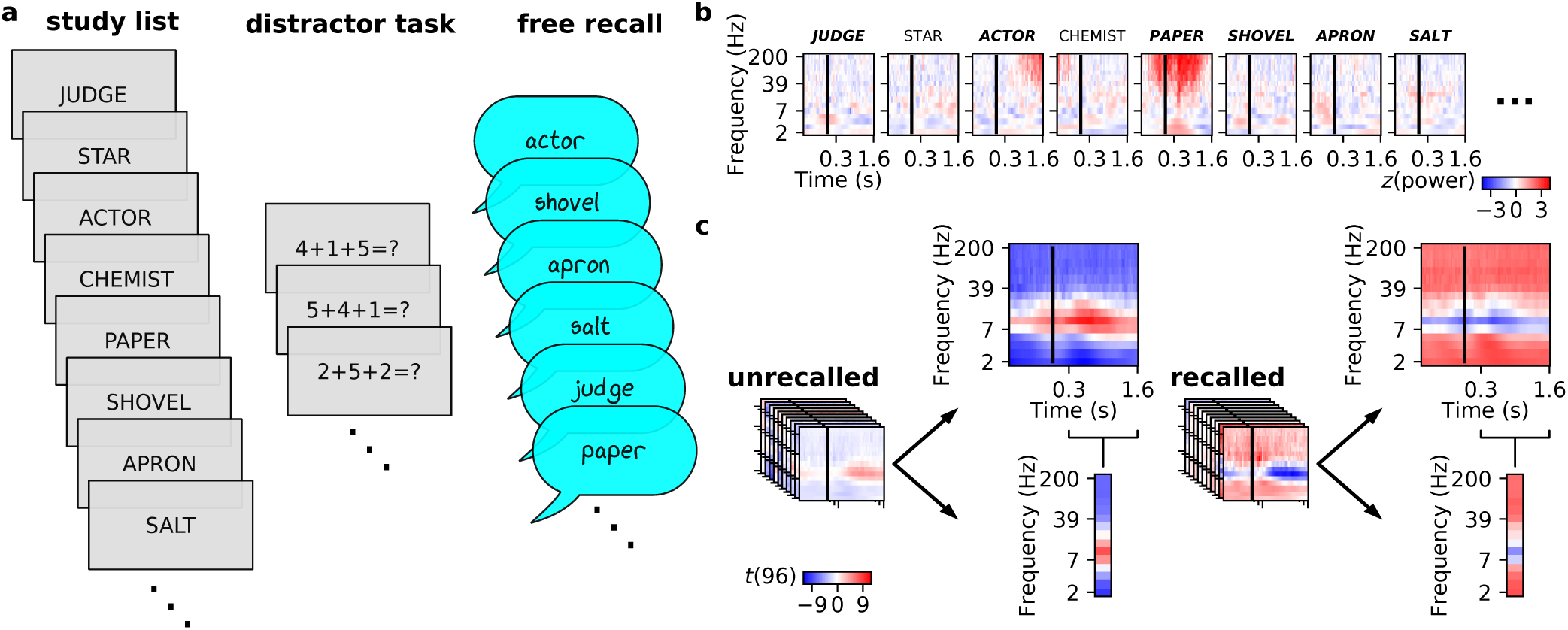
(**a**) Illustration of an individual trial in our experiment consisting of a study list followed by a distractor task, and a free recall test. There were 24 of these trials in each experimental session and each study list consisted of 24 items. See methods for details. (**b**) *z*-transformed power around the presentation of study words during the beginning and end of one participant’s (ID: 374) 4th study list in the 16th experimental session. The study words are indicated at the top of each subpanel with bold italic font indicating subsequent recall. (**c**) Average power for subsequently unrecalled (left) and subsequently recalled (right) words during study across all lists from all participants (we averaged all data within participants and calculated the shown *t*-values across participants). All of our analyses were based on neural activity between 0.3 and 1.6 s following study word onset (indicated with vertical black lines) and the average power across this time interval is also illustrated. For this visualization, we aggregated EEG activity across 28 superior electrodes (see methods for details).

For our second approach we calculated list-level SMEs (rather than the standard item-level SMEs), leveraging evidence that endogenous factors associated with cognitive function vary slowly. Specifically, sequential dependencies in human performance as well as investigations of endogenous neural fluctuations that drive variability in evoked brain activity and overt behavior suggest that endogenous factors operate at time scales that are slower than the time allocated to the study of individual items in standard memory tasks (many seconds or minutes rather than a few seconds or less; Kahana et al., 2018; Gilden, Thornton, & Mallon, 1995; Mueller & Weidemann, 2008; Verplanck, Collier, & Cotton, 1952; Monto, Palva, Voipio, & Palva, 2008; Schroeder & Lakatos, 2009; Arieli, Sterkin, Grinvald, & Aertsen, 1996; Fox, Snyder, Zacks, & Raichle, 2005; Fox, Snyder, Vincent, & Raichle, 2007; Fox & Raichle, 2007; Raichle, 2015). To calculate list-level SMEs, we averaged epochs of EEG activity following the presentation of individual study items within each list and used these list-averaged epochs to predict the proportion of recalled words in each list. This approach eliminates or severely reduces the effects of item-specific external factors (because we are averaging neural activity across all study periods in a list), but the list-level SME could still reflect other external factors that also affect recall performance (such as session-level time-of-day effects or list-level proactive interference effects; Kahana et al., 2018). We therefore also statistically removed effects of list and session number (as well as effects of the average “recallability” of the words comprising each list; see methods for details) and computed SMEs for the corresponding residuals. As with the item-level SMEs, these residuals highlight contributions of endogenous factors. Comparing the SMEs for list-level recall to the SMEs for residuals of list-level recall after accounting for external factors associated with each list and experimental session thus allowed us to estimate the extent to which list-level SMEs are driven by endogenous factors associated with encoding success.

## Methods

### Participants

We analyzed data from 97 young adults (18–35) who completed at least 20 sessions in Experiment 4 of the Penn Electrophysiology of Encoding and Retrieval Study (PEERS) in exchange for monetary compensation. This study was approved by the Institutional Review Board at the University of Pennsylvania and we obtained informed consent from all participants. Recall performance for a large subset of the current data set was previously reported (Kahana et al., 2018), but this is the first report of electrophysiological data from this experiment. Data from PEERS experiments are freely available at http://memory.psych.upenn.edu and have been reported in several previous publications (Healey, Crutchley, & Kahana, 2014; Healey & Kahana, 2014, 2018; Lohnas & Kahana, 2013; Siegel & Kahana, 2014; Lohnas, Polyn, & Kahana, 2015; Weidemann & Kahana, 2016, 2019). Our analyses included data from all participants with at least 20 sessions.

### Experimental task

Each of up to 23 experimental sessions consisted of 24 study lists that each were followed by a delayed free recall test. Specifically, each study list presented 24 session-unique English words sequentially for 1,600 ms each with a blank inter-stimulus interval that was randomly jittered (following a uniform distribution) between 800 and 1,200 ms. After the last word in each list, participants were asked to solve a series of arithmetic problems of the form *A* + *B* + *C* =? where, *A, B*, and *C* were integers in [1, 9]. Participants responded to each problem by typing the result and were rewarded with a monetary bonus for each correctly solved equation. These arithmetic problems were displayed until 24 s had elapsed and were then followed by a blank screen randomly jittered (following a uniform distribution) to last between 1,200 and 1,400 ms. Following this delay, a row of asterisks and a tone signaled the beginning of a 75 s free recall period. A random half of the study lists (except for the first list in each session) were also preceded by the same arithmetic distractor task which was separated from the first study-item presentation by a random delay jittered (following a uniform distribution) to last between 800 and 1,200 ms. Each session was partitioned into 3 blocks of 8 lists each and blocks were separated by short (approximately 5 min) breaks. At each session participants were asked to rate their alertness and indicate the number of hours they had slept in the previous night.

### Stimuli

Across all lists in each session the same 576 common English words (24 words in each of 24 lists) were presented for study, but their arrangement into lists differed from session to session (subject to constraints on semantic similarity; Healey et al., 2014). These 576 words were selected from a larger word pool (comprising 1,638 words) used in other PEERS experiments. The 576-word subset of this pool used in the current experiment is included as supplementary material and ranged in arousal (2.24–7.45, *M* = 4.04) and valence (1.71–8.05, *M* = 5.52) according to independent ratings on these dimensions on scales between 1 and 9 (Warriner, Kuperman, & Brysbaert, 2013). Many participants also returned for a 24th session that used words from the entire 1,638-word pool, but we are not reporting data from that session here. We estimated the mean recallability of items in a list from the proportion of times each word within the list was recalled by other participants in this study.

### EEG data collection and processing

Electroencephalogram (EEG) data were recorded with either a 129 channel Geodesic Sensor net using the Netstation acquisition environment (Electrical Geodesics, Inc.; EGI) or with a 128 channel Biosemi Active Two system. EEG recordings were re-referenced offline to the average reference. Because our regression models weighted neural features with respect to their ability to predict (residuals of) recall performance in held out sessions, we did not try to separately eliminate artifacts in our EEG data. Data from each participant were recorded with the same EEG system throughout all sessions and for those sessions recorded with the Geodesic Sensor net, we excluded 26 electrodes that were placed on the face and neck, rather than the scalp, from further analyses. For the visualization of EEG activity in the figures, we aggregated over electrodes 4, 5, 12, 13, 19, 20, 24, 28, 29, 37, 42, 52, 53, 54, 60, 61, 78, 79, 85, 86, 87, 92, 93, 111, 112, 117, 118, and 124 for the EGI system and electrodes A5, A6, A7, A18, A31, A32, B2, B3, B4, B18, B19, B31, B32, C2, C3, C4, C11, C12, C24, C25, D2, D3, D4, D12, D13, D16, D17, and D28 for the Biosemi system. These correspond to the superior regions of interest we used previously (Weidemann, Mollison, & Kahana, 2009). All of our classification and regression models, however, used measures from all individual electrodes (with the exception of those covering the face and neck for the EGI system) as input without any averaging across electrodes. The EGI system recorded data with a 0.1 Hz high-pass filter and we applied a corresponding high-pass filter to the data collected with the Biosemi system. We used MNE (Gramfort et al., 2013, 2014), the Python Time-Series Analysis (PTSA) library (https://github.com/pennmem/ptsa_new), Sklearn (Pedregosa et al., 2011) and custom code for all analyses.

We first partitioned EEG data into epochs starting 800 ms before the onset of each word in the study lists and ending with its offset (i.e., 1,600 ms after word onset). We also included an additional 1,200 ms buffer on each end of each epoch to eliminate edge effects in the wavelet transform. We calculated power in 15 logarithmically spaced frequencies between 2 and 200 Hz, applied a log-transform, and down-sampled the resulting time series of log-power values to 50 Hz. We then truncated each epoch to 300–1,600 ms after word onset. For the item-based models we used each item’s *z*-transformed mean power in each frequency across this 1,300 ms interval as features to predict (residual) subsequent recall. For the list-based regression models we averaged these values across all items in each list to predict (residuals of) list-level recall.

### Removing effects of external factors

For the item based analyses we fit logistic regression models separately for each participant to predict each item’s recall from its average recallability (i.e., it’s average probability of recall calculated from all other participants’ recall data), its serial position within the study list, the list number within the current session, and the session number within the experiment. We treated all of these predictors, except for recallability, as categorical to accommodate any functional relationship between them and recall performance. This allowed us to use list and session number as predictors to model the combined effects of list and session-specific external factors rather than attempting to capture each of them separately. Furthermore, fitting these models separately to each participant’s data allowed us to accommodate potentially idiosyncratic relationships between external factors and the predictors in our model as well as those between external factors and recall performance. We then calculated residuals from the full model including all item-level predictors as well as from nested models including all but one of the predictors as described in the main text. Residuals from logistic regression models are constrained to fall between −1 and 1 (assuming the two possible outcomes are coded as 0 and 1). To make these residuals more similar to those from the linear regression models, we transformed the residuals to fall between 0 and 1 (just like list-level recall probabilities) and then applied a logit-transform: 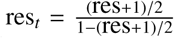, where res_*t*_ and res are the transformed and untransformed residuals respectively. All references to residuals from logistic regression models in other parts of this paper refer to transformed residuals.

For the list-based analyses we proceeded similarly, fitting linear regression models separately for each participant to predict the logit transformed probability of recall for each list (i.e., the proportion of words that were recalled in each list). We used the average recallability of words within each list, list number within each session, and session number within the experiment as predictors (treating list and session number as categorical predictors). We again calculated residuals for the full model and also for two nested models: one including average recallability for each list and list number (list-level predictors) and one only including session number (session-level predictor).

### Item-based classifier

For the item-based classifier we used a nested crossvalidation procedure to simultaneously determine the regularization parameter and performance of L2-regularized logistic regression models predicting each item’s subsequent recall. We applied this nested cross-validation approach separately to the data from each participant to accommodate idiosyncratic relationships between brain activity and recall performance and inter-individual differences in signal quality. At the top level of the nested cross-validation procedure we held out each session once—these held out sessions were used to assess the performance of the models. Within the remaining sessions, we again held out each session once— these held-out sessions from within each top-level cross-validation fold were used to determine the optimal regularization parameter, *C*, for Sklearn’s LogisticRegression class. We fit models with 9 different *C* values between 0.00002 and 1 to the remaining sessions within each cross-validation fold and evaluated their performance as a function of *C* on the basis of the held out sessions within this fold. We then fit another logistic regression model using the best-performing *C* value to all sessions within each cross-validation fold and determined the model predictions on the sessions that were held-out at the top level. We determined the performance of our models solely on the basis of the predictions from these held-out sessions. There are many reasonable alternatives for setting up these models; our choice of L2 regularization was motivated by good performance of these models in similar data sets (Weidemann & Kahana, 2019; Weidemann et al., 2019), and not informed by the current results.

### Item and list-based regression models

For the item- and list-based regression models we followed the same procedure as for the item-based classifier to determine the optimal level of regularization for L2 regularized linear regression models predicting residuals of item-level recall or (residuals of) list-level recall performance. Specifically, we used the same nested cross-validation procedure described above to determine optimal values for α (corresponding to 1/*C*), the regularization parameter in Sklearn’s Ridge class, testing 9 values between 1 and 65536. We applied these models to the (logit-transformed) proportion of items recalled for each list and to the residuals from the various item- and list-level models as described above. Thus, in cases where we investigated the effects of external factors on recall performance, we first fit a regression model predicting recall performance on the basis of the external factor(s) (as described above in the “Removing effects of external factors” section) and then used a ridge regression model to predict the residuals from that model fit (“residual recall performance”) on the basis of brain activity.

### Shuffled control lists

For our list-level analyses we also computed SMEs for shuffled control lists to investigate the extent to which SMEs were linked to individual item properties or instead relied on slowly varying endogenous factors. If the list-level SME is merely an average of individual item level SMEs, shuffling individual items (together with the corresponding neural activity) should not affect the list-level SME as long as the shuffled lists contain the same number of subsequently recalled and not subsequently recalled items. Effects of slowly varying endogenous factors would, however, be disrupted if items and associated neural activity were rearranged. For this approach, we separated all recalled and unrecalled items (together with the corresponding neural activity) in each session, shuffled both sets of items separately (keeping each item linked with its neural activity), and then synthesized new lists with the original proportions of recalled and unrecalled items from the shuffled pools of recalled and unrecalled items. We repeated this procedure 20 times for each participant and concatenated the resulting shuffled lists. This shuffled session thus consisted of 20 copies of each item synthesized into 480 lists that matched the recall performance of the 24 original lists (the performance of each original list was represented 20 times in the shuffled session). We then applied all of our list-level SME analyses to these shuffled lists. Specifically, we predicted (residuals of) list-level recall performance on the basis of the neural data associated with the items making up the synthesized lists.

## Results

The standard item-level subsequent memory analysis contrasts neural activity during the encoding of subsequently recalled and non-recalled items. The present experiment sequentially displayed lists of items (words) for study and tested memory in a delayed free recall task (Figure 1a). During the encoding period of each studied item, we calculated the spectral power of the EEG signal at frequencies between 2 and 200 Hz. Figure 1b shows an excerpt of an actual study list with associated *z*-transformed spectral power, shown as a joint function of encoding time and frequency for each excerpted item. The average time-frequency spectrogram for recalled and non-recalled items, shown in Figure 1c, illustrates the spectral subsequent memory effect reported in prior studies (Paller & Wagner, 2002; Hanslmayr & Staudigl, 2014). Specifically, subsequently recalled items exhibit greater high frequency (> 30 Hz) activity and reduced alpha power (8–12 Hz) as compared with not-recalled items. Before commencing our analyses we had decided to focus on a time window between 0.3 and 1.6 s following the onset of each study item to maximize our chance of capturing item-specific effects in our SME contrasts. However, as is evident in Figure 1c, the SME was sustained throughout the entire 1.6 s during which the item appeared on the screen and also in the pre-stimulus interval (consistent with previous reports of pre-stimulus SMEs; Sweeney-Reed et al., 2016; Otten, Quayle, Akram, Ditewig, & Rugg, 2006; Fellner et al., 2013; Guderian, Schott, Richardson-Klavehn, & Duzel, 2009; Park & Rugg, 2010; Urgolites et al., 2020).

The power of the SME analysis lies in its ability to reveal encoding processes that lead to successful recall. However, the standard item-level SME conflates a multitude of factors that determine the recallability of any given item. The position of an item in the study list constitutes one such factor. The top of Figure 2 illustrates the serial position effect in our delayed free-recall experiment. As expected based on prior work, we observed superior recall for early list items (the so-called primacy effect). The mental arithmetic task between study and test attenuates the recency effect that is typical of immediate recall (Murdock, 1962). Given the strong effect of serial position on recall performance, we can expect any SME to also reflect a contrast of neural activity associated with different serial positions. The second row of Figure 2 shows the neural activity associated with the encoding interval at each serial position irrespective of recall status. Here one sees a marked shift in neural activity across serial positions: Neural activity at early serial positions resembles that associated with recalled items and that at later serial positions is similar to that associated with not-recalled items (cf. Figure 1c). The last two rows of Figure 2 illustrate that this pattern is not simply due to the confound between recalled status and serial position: Even when we plot the pattern of spectral activity as a function of serial position separately for recalled and not-recalled items, neural activity at early serial positions resembles that associated with recalled items and that at later serial positions is more similar to that associated with not-recalled items in the standard SME (cf. Figure 1c). This illustrates how the subsequent memory analysis can be misleading: differences between recalled and non-recalled items may be indexing differences between primacy and non-primacy items. Controlling for the effect of serial position represents a logical solution to this problem. However, serial position is but one of many variables known to influence recall performance. We thus introduce a statistical framework to separate the effects of known external factors from the hypothesized endogenous variability driving encoding success, as described below.

**Figure 2.**
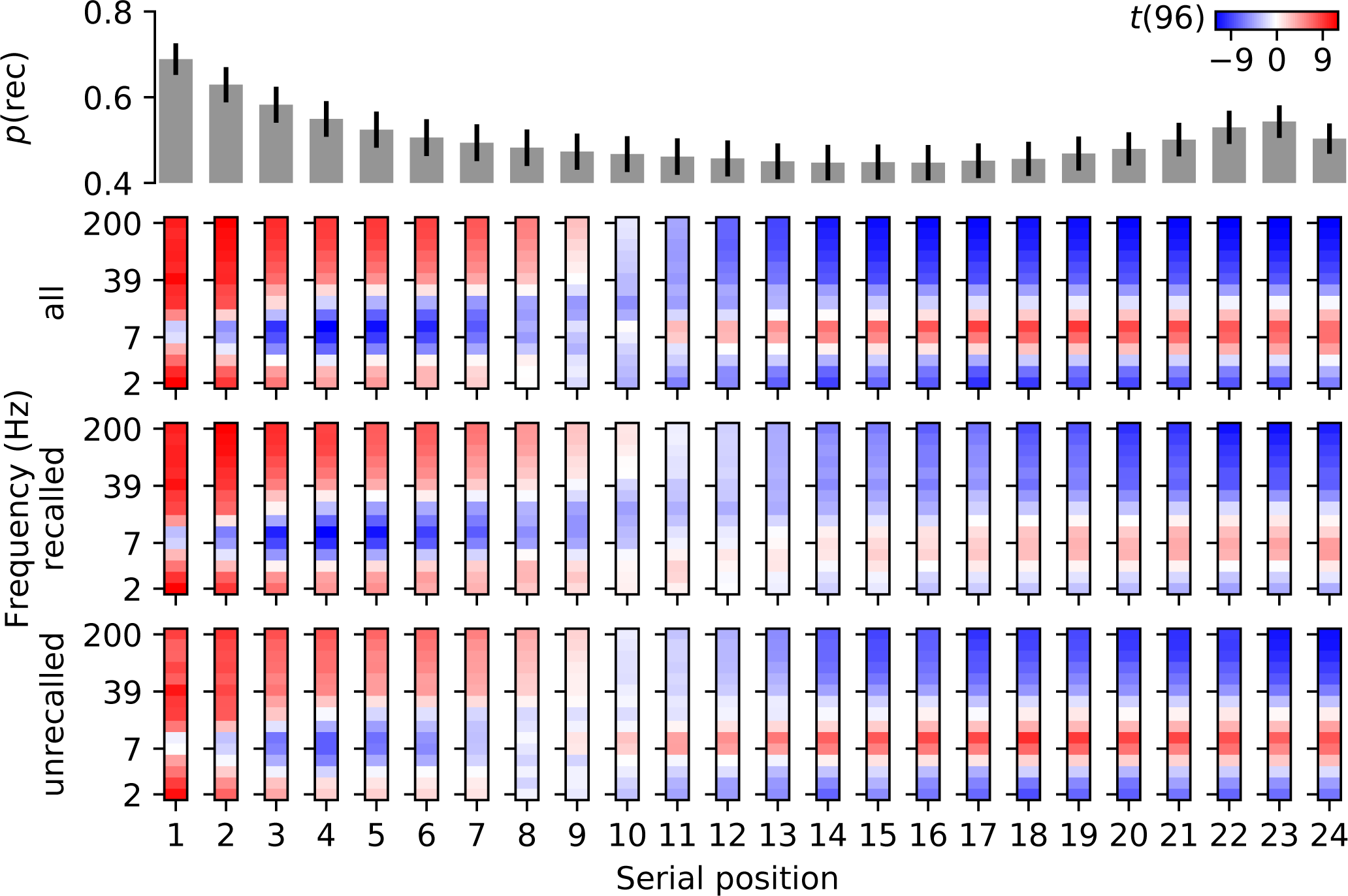
Mean probability of recall as a function of serial position across all participants (top row) and associated neural activity (averaged between 0.3 and 1.6 s after the onset of study items) for all, subsequently recalled, and subsequently not-recalled trials respectively (we averaged all data within participants and calculated the shown *t*-values across participants). Error bars indicate 95% confidence intervals. For this visualization, we aggregated EEG activity across 28 superior electrodes (see methods for details).

Our analytic approach combines multivariate classification of neural data (Ezzyat et al., 2018; Weidemann et al., 2019; Weidemann & Kahana, 2019) with a multi-factor model of external variables shown to influence item-level recall performance (Kahana et al., 2018). To implement a multivariate analogue to the standard SME analysis, we trained L2 regularized logistic regression classifiers using brain activity to predict the recall status of individual items (the performance of these models indexes what we refer to as an “uncorrected SME”). We also trained L2 regularized linear regression models using brain activity to predict residuals of recall performance after statistically controlling for the effects of external factors that also predict recall performance (the performance of these models indexes what we refer to as a “corrected SME”).

For both uncorrected and corrected SMEs, we evaluate how well each model predicts (residuals of) recall performance in held out sessions. Typical metrics of model performance differ between binary classification (as in our uncorrected SME analyses) and continuous regression models (as in our corrected SME analyses). To directly compare both types of SMEs, we computed correlations between model predictions and (residual) recall performance. For the uncorrected SME, this is a point-biserial correlation because recall performance is a binary variable (each item is either recalled or not) and the model prediction is a continuous measure corresponding to the predicted recall probability of each item. For the corrected SME, this is a standard product-moment correlation between the continuous residual recall performance and the continuous model prediction (see *Methods* for details). Both of these models use spectral features of EEG activity during word encoding to predict that item’s (residual) recall status.

The correlation between model predictions and (residual) item-level recall performance quantifies the association between neural features during encoding and subsequent (residual) recall performance—it serves as our multivariate SME measure. The top of Figure 3a shows the distribution of these correlations across participants for the uncorrected SME (distribution marked “item”) relating neural features to the recalled status of individual items. This uncorrected SME was significant (*M* = 0.16, *t*(96) = 22.681, *SE* = 0.007, *p* < 0.001, *d* = 2.303) indicating that the different average activity patterns for recalled and not-recalled items shown in Figure 1c were indeed associated with a reliable item-level SME. The next distribution (labeled “item|all”) corresponds to the corrected SME statistically controlling for all external factors. Specifically, these correlations quantify the relation between neural features and the residuals of logistic regression models predicting recall status on the basis of individual item-recallability, serial position, list number within the current session, and session number within the experiment. This corrected SME, was also statistically significant (*M* = 0.12, *t*(96) = 19.015, *SE* = 0.006, *p* < 0.001, *d* = 1.931), indicating a substantial SME, even after controlling for external factors. The size of this SME was somewhat smaller than that for the uncorrected recall performance (*t*(96) = 9.738, *SE* = 0.004, *p* < 0.001, *d* = 0.989) reflecting the fact that the uncorrected SME does include the effects of some external factors.

**Figure 3.**
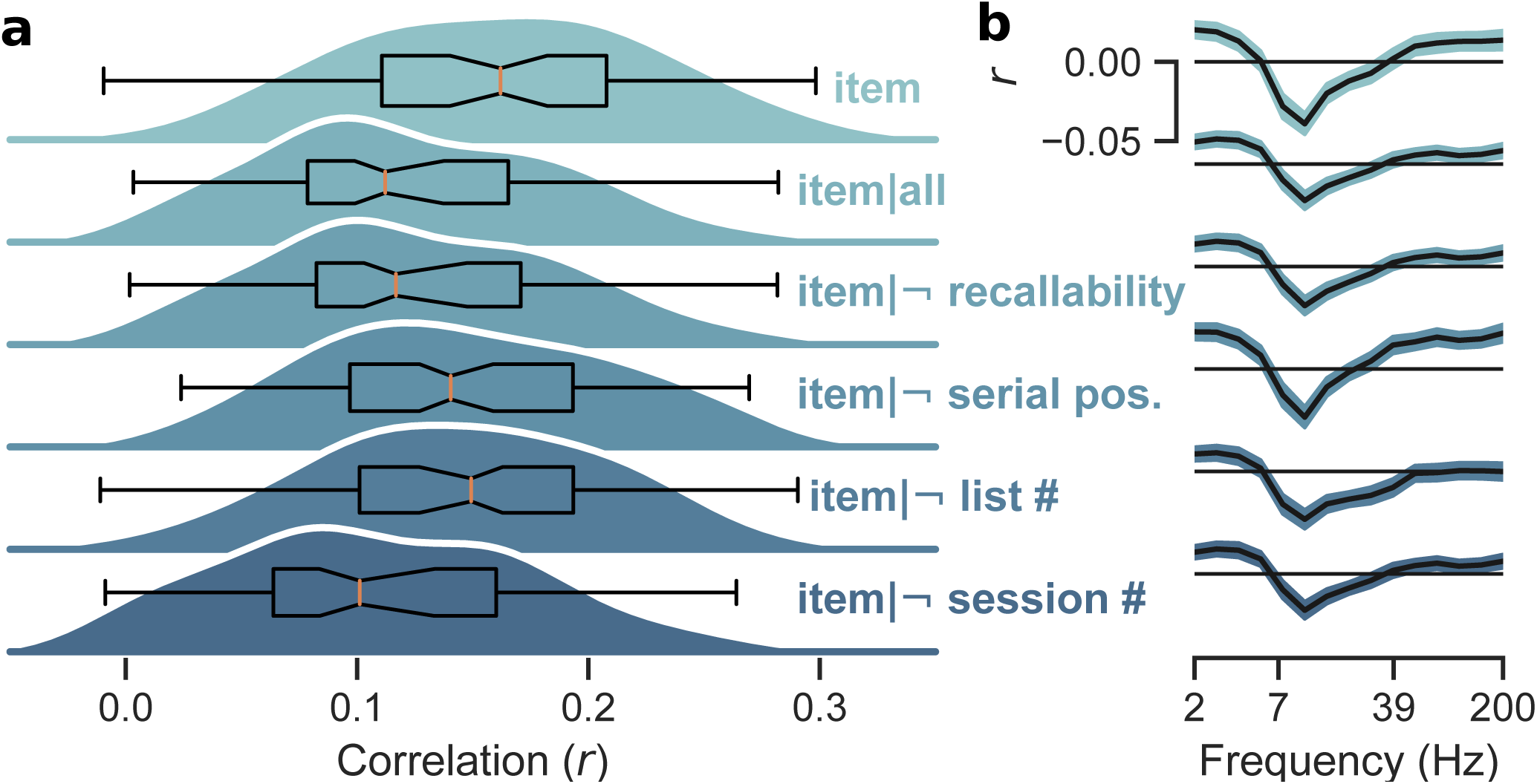
(**a**) Distribution of uncorrected item-level SMEs (“item”) across all participants and of corresponding corrected SMEs accounting for all factors or all but the indicated factor respectively (a prefix signifies that the indicated factor was omitted). Overlaid boxplots indicate the quartiles of the distribution with a notch showing the bootstrapped 95% CI around the median. Whiskers extend to 1.5× the inter-quartile range. (**b**) Mean correlations between power at different frequencies (aggregated across 28 superior electrodes) and the respective (residuals of) item-level recall performance across all participants (lined up with the corresponding SMEs in Panel **a**). The black horizontal lines indicate zero. Error regions indicate 95% CIs.

To better understand how the different factors affect the SME, we repeated this analysis, but held out each of the external factors in turn. Specifically we computed four partially-corrected SMEs that each omitted one external factor. The remaining parts of Figure 3a show the results of these analyses without controlling for the effects of recallability, serial position, list number, and session number respectively. All resulting SMEs are positive (*M* = 0.11–0.15, *t*(96) = 16.341–22.471, *SE* = 0.006–0.007, *p*s < 0.001, *d* = 1.659–2.282) and significantly different from the SME for uncorrected recall performance (*t*(96) = 4.726–13.438, *SE* = 0.003–0.004, *p*s < 0.001, *d* = 0.479–1.364) as well as from that correcting for all external factors (*t*(96) = 5.939–10.790, *SE* = 0.001–0.003, *p*s < 0.001, *d* = 0.603–1.096). This indicates that each of the external factors contributes to the difference between the size of the uncorrected and the corrected SME and that none of these factors can account for this difference in isolation. Serial position, however, explains most of this discrepancy—when controlling for all other factors, the corresponding SME is almost as large as the uncorrected SME (mean correlation of 0.15 as opposed to 0.16) and additionally also controlling for serial position is responsible for reducing the SME to a mean correlation of 0.12.

To the extent that the uncorrected SME reflects both endogenous and external factors, we would expect that statistically removing the effects of external factors would reduce the size of the SME. Correspondingly, only partially removing effects of external factors (e.g., by holding out the removal of one of the external factors like we did in the analyses described above) should result in SMEs that fall somewhere between the uncorrected SME and the SME correcting for more external factors. This is the pattern we observed, with one exception: when we statistically removed the effects of all factors except for the session number, the resulting SME was slightly smaller than that for the SME also removing that effect (mean correlation of 0.11 as opposed to 0.12). This indicates that recall performance varies with session number, but that this effect of session number is not effectively captured by our measures of brain activity. Hence, when we statistically controlled for the effects of session number we removed variability in recall performance that we could not account for with our measures of brain activity, leading to a slightly larger SME (and, conversely, a failure to remove the effects of session number reduced the SME).

As Figure 3a also shows, there was substantial overlap between the distributions for the uncorrected and corrected SMEs demonstrating that the effects of external factors were small relative to the size of the SME. Specifically, the effect sizes associated with the uncorrected and corrected SMEs corresponded to Cohen’s (Cohen, 1988) *d*s of 2.303 and 1.931, respectively (with the Cohen’s *d*s for corrected SMEs holding out one of the factors ranging between 1.659 and 2.282). The difference between the uncorrected and corrected SME was about half that size (Cohen’s *d* of 0.989 and 0.479–1.364 for the differences between the uncorrected SME and the corrected SMEs holding out one of the factors). Another way to interpret the sizes of the uncorrected and corrected SMEs relative to their difference is by directly evaluating the corresponding correlations and their difference. According to Cohen’s convention, the correlations for all SMEs correspond to a small effect size (0.1 < *r* < 0.3). Differences in correlations can be assessed with Cohen’s *q* (i.e., the difference between the Fisher-*z* transformed correlations) which is 0.041 for the difference between the uncorrected and corrected SME (and ranges between 0.018 and 0.054 for the differences between the uncorrected SME and the corrected SME holding out one of the factors)—all well below the threshold Cohen proposed for a small effect (0.1 < *q* < 0.3).

Figure 3b shows correlations between power at different frequencies and (residual) recall performance to help illustrate the importance of different features for our regularized logistic and linear regression models relating brain activity to (residual) recall performance. Across all measures of (residual) recall performance, correlations with spectral power were more negative in the α range (around 10 Hz) and less negative at higher and lower frequencies. The correlations between power and uncorrected item-level recall were positive for frequencies in the γ range (> 40 Hz)—an effect that was substantially reduced for all item-level residuals, except for that not correcting for serial position. This suggests that positive correlations between γ power and recall performance largely reflect serial position effects (see also Figure 2).

Rather than statistically controlling for factors that were specific to individual items (i.e., serial position and recallability), our list-level SME eliminates or severely reduces these factors by averaging brain activity over the encoding epochs to predict (residuals of) the proportion of recalled items in each list. Because each list contained the same number of items, effects of serial position averaged out, eliminating this factor from affecting list-level SMEs. Even though recallability is specific to individual items, lists could vary with respect to the average recallability of their constituent items. We therefore considered not only list number and session number, but also average recallability of items within the list as external factors to control for in our calculation of corrected list-level SMEs. As for our item-level SMEs, we quantify list-level SMEs by calculating the correlations between predictions from L2 regularized linear regression models and (residual) recall performance.

The top of Figure 4a (labeled “list”) shows the distribution of the uncorrected list-level SME (*M* = 0.26, *t*(96) = 18.213, *SE* = 0.015, *p* < 0.001, *d* = 1.849). It is tempting to compare the size of this list-level SME to the item-level SME shown at the top row of Figure 3a, but such direct comparisons are difficult to make sensibly. The EEG features driving the list-level SME were averaged across all study epochs within each list, whereas the item-level SME relied on features from individual epochs. Thus the neural features making up the item and list-level SMEs may differ substantially in their respective signal to noise ratios and the number of observations contributing to these different kinds of SMEs also differed considerably (in our case by a factor of 24, because each list consisted of 24 items).

**Figure 4.**
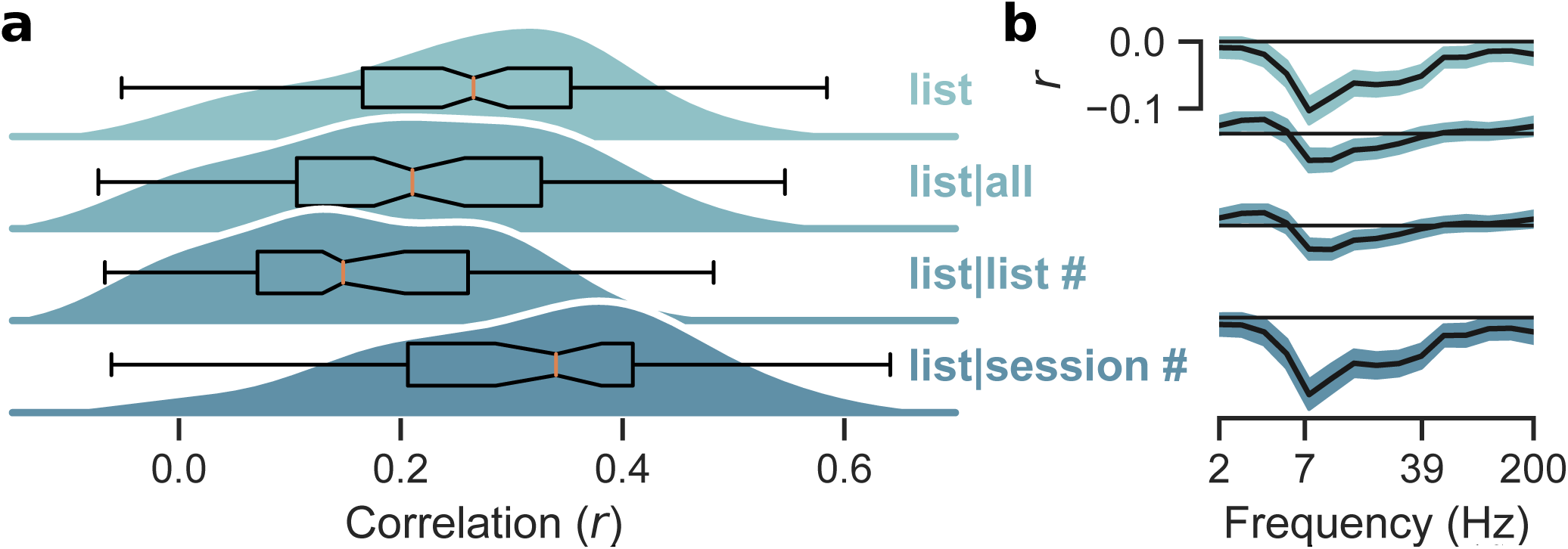
(**a**) Distribution of uncorrected list-level SMEs (“list”) across all participants and of corresponding corrected SMEs accounting for all factors or only the indicated ones (here “list #” refers to the joint effects of both list number and average recallability of words in each list). Boxplots are as in Figure 3. (**b**) Mean correlations between power at different frequencies (aggregated across 28 superior electrodes) and the respective (residuals of) list-level recall performance across all participants (lined up with the corresponding SMEs in Panel **a**). The black horizontal lines indicate zero. Error regions indicate 95% CIs.

To calculate corrected list-level SMEs, we fit linear regression models to predict list-level recall performance on the basis of average recallability of items in that list, list number, and session number. We then used brain activity to predict residual list-level recall performance. The second row of Figure 4a (labeled “list|all”) shows this corrected list-level SMEs (*M* = 0.22, *t*(96) = 14.332, *SE* = 0.015, *p* < 0.001, *d* = 1.455). This effect was smaller than the uncorrected list-level SME (*t*(96) = 5.548, *SE* = 0.008, *p* < 0.001, *d* = 0.563), reflecting the fact that external factors do contribute to the uncorrected list-level SME. The fact that we could demonstrate a sizable corrected list-level SME, however, supports our previous result that external factors are not critical drivers of the SME.

To better understand the extent to which list and session-level external factors contribute to the list-level SME, we statistically controlled for average recallability of items within each list and list number (list-level effects; third row of Figure 4b labeled “list|list #”) and, separately, for session number (session-level effects; fourth row of Figure 4b labeled “list|session #”). The corresponding SMEs were significant (*M* = 0.16 and 0.32, *t*(96) = 12.668 and 20.132, *SE* = 0.013 and 0.016, respectively, both *p*s < 0.001, *d* = 1.286 and 2.044, respectively). Their sizes, however, fell outside the range spanned by the SME controlling for all external factors and the uncorrected SME. The SME correcting for list-level factors was smaller than that correcting for all external factors and the uncorrected SME (*t*(96) = 11.606 and 12.466, *SE* = 0.005 and 0.008, respectively, both *p*s < 0.001, *d* = 1.178 and 1.266, respectively), whereas the SME correcting for session was larger than both (*t*(96) = 13.134 and 13.950, *SE* = 0.009 and 0.005, respectively, both *p*s < 0.001, *d* = 1.333 and 1.416, respectively). This pattern confirms our previous finding that our measures of brain activity did not effectively capture session-level external factors that affect recall performance. Hence, statistically controlling for their effects enhances our ability to predict residual recall performance from brain activity whereas a failure to remove that variability from recall performance reduces the SME.

As for the item-level SMEs, Figure 4a shows substantial overlap between the distributions for the uncorrected and corrected list-level SMEs. Analyses of corresponding effect sizes confirm that here, too, effects of external factors were small relative to the size of the SME. Specifically Cohen’s *d* for the uncorrected and corrected SMEs were 1.849 and 1.455, respectively (corresponding *d*s for the corrected SME considering only list or session-related factors were 1.286 and 2.044 respectively). The size of the difference between the uncorrected and the corrected SME was only about a third (*d* = 0.563) of the individual effects (but, *d* = 1.266 and 1.416 for the corrected SMEs only accounting for list and session-related factors, respectively). As before, we can also interpret the size of these effects by considering the corresponding correlations directly. From that perspective, the uncorrected and all corrected SMEs correspond to small effects (0.1 < *r* < 0.3) whereas the differences between the uncorrected and the corrected SME falls short of a small effect (*q* = 0.047; corresponding *q*s for the differences with corrected SMEs considering only list or session-related factors were 0.1 and 0.07 respectively).

Just as in Figure 3b, Figure 4b shows the correlations between power in different frequencies and (residuals of) recall performance. The qualitative pattern of these correlations aligned with the pattern for item-level SMEs with more negative correlations in the α range and less negative correlations at lower and higher frequencies. Positive correlations between γ power and (residuals of) list-level recall performance were absent, supporting our previous interpretation that these positive correlations in item-level SMEs are largely driven by serial position effects (which are averaged out in the list-level analyses).

The presence of a robust list-level SME is compatible with endogenous factors that vary slowly (over many seconds or minutes) rather than with the presentation of individual items during the study list. Indeed, to the extent that factors driving the SME are closely linked to the presentation of individual items, characterizing these factors as “endogenous” would be problematic. To investigate the extent to which factors predicting subsequent recall are tied to individual items rather than varying more slowly over the study periods we constructed shuffled lists that mirrored the distribution of recall performance, but synthesized lists from randomly selected items within each session. Figure 5 shows the list-level SMEs for these shuffled lists. As is evident from the Figure, this shuffling procedure practically eliminated the SME. High statistical power resulted in statistically significant deviations from zero, but the largest shuffled SME corresponded to a mean correlation of 0.03 with the residual recall performance after accounting for session effects which was an order of magnitude smaller than the corresponding unshuffled SME. All shuffled SMEs were significantly smaller than the corresponding unshuffled ones (*t*(96) = 14.286–20.361, *SE* = 0.013–0.016, *p*s < 0.001, *d* = 1.450–2.067), supporting our previous result that (slowly varying) endogenous factors (rather than item-specific, or otherwise external, factors) are the main drivers of the SME.

**Figure 5.**
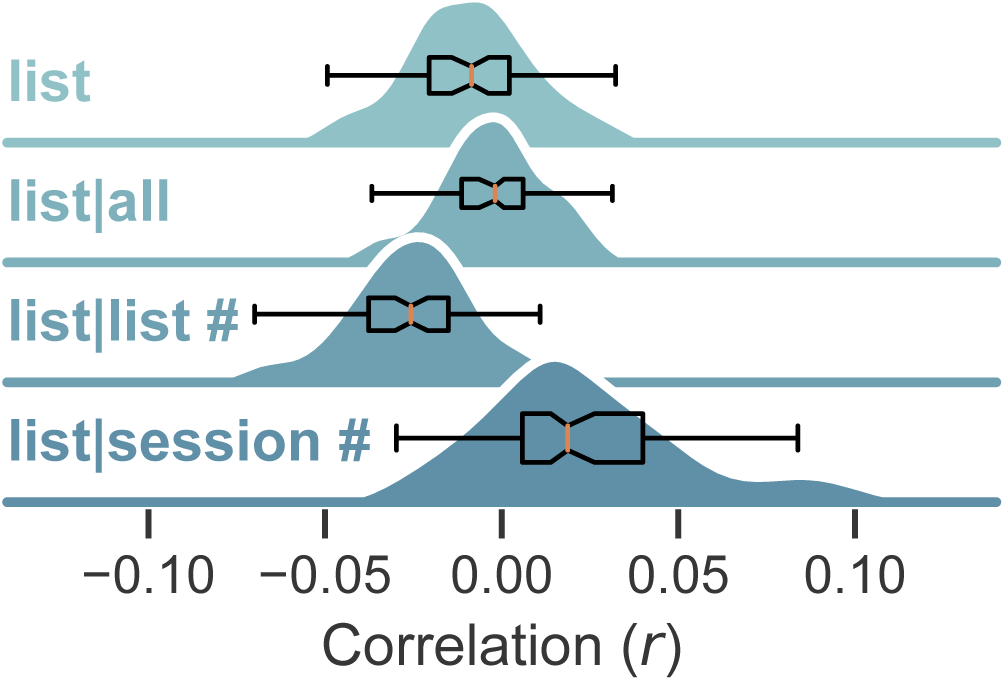
Distribution of uncorrected list-level SMEs (“list”) across all participants for synthesized lists made up from randomly selected items within a session (see methods for details) and of corresponding corrected SMEs accounting for all factors or only the indicated ones (here “list #” refers to the joint effects of both list number and average recallability of words in each list). Boxplots are as in Figures 3 and 4.

## Discussion

The subsequent memory analysis of neural data has provided researchers with a powerful tool for uncovering the brain mechanisms that underlie successful memory formation. Armed with this methodology, cognitive neuroscientists have conducted hundreds of experiments, using a wide range of recording techniques, seeking to elucidate the brain signals and networks that accompany memory acquisition. Yet, despite an impressive body of data amassed in recent decades, key questions about the neural correlates of memory acquisition remain unanswered. Specifically, to what extent do these neural correlates reflect known external factors that determine memorability, or endogenously varying brain states that determine the efficiency of memory acquisition? Prior research suggests that both external and endogenous factors play a role: On the one hand, experimental manipulations of item encoding affect the SME (Otten & Rugg, 2001; Staudigl & Hanslmayr, 2013; Fellner et al., 2013), suggesting a role for external factors. On the other hand, neural activity prior to item onset predicts subsequent memory, suggesting a role for endogenous factors unrelated to item processing (Sweeney-Reed et al., 2016; Otten et al., 2006; Fellner et al., 2013; Guderian et al., 2009; Park & Rugg, 2010; Urgolites et al., 2020). We approached this question by examining how the SME changed after statistically controlling for a comprehensive set of external factors. We also sought to remove effects of item-specific external factors by aggregating brain activity over the study periods of all items within a list to predict list-level recall (i.e., a list-level SME). Both approaches for removing the effects of external factors resulted in relatively modest decreases to the SME, implicating endogenous factors as the main drivers of the SME.

Because it is impossible to perfectly control for effects of all possible external factors, distinguishing between effects of external variables and endogenous processes is notoriously difficult. We approached this challenge by treating serial position, list, and session number as categorical predictors, effectively modeling the joint effects of external factors associated with these predictors without having to commit to a particular functional form relating these predictors to recall performance. By fitting these models separately to the data from each individual, we were also able to accommodate individual differences. Our approach attributed any variability in recall performance that covaried with one of our external factors to that factor, even though it is likely that some of that variability could reasonably be classified as “endogenous” (e.g., sessions could be administered at different times from day to day, and corresponding effects of circadian rhythms would have been classified as an external session effect). Because of the fact that our external factors likely represented the joint effects (including interactions) of a large number of factors, we did not explicitly model any interactions between the factors we considered. Such interactions would be difficult to interpret and we would expect them to be small given that they would reflect consistent relationships between somewhat arbitrary groups of of factors. Our approach to modeling external factors thus should yield a conservative estimate of the contributions of endogenous factors, despite the fact that we cannot completely rule out contributions of external factors (and some corresponding interactions) to our corrected SMEs.

Our findings of strong list-level SMEs, and their elimination when synthesizing lists of randomly selected items within a session, provide strong additional evidence against the interpretation that the SME reflects item-level factors that influence memorability. Specifically, the elimination of the list-level SME for shuffled lists shows that the list-level SME is not simply an aggregation of neural activity predicting recall success for individual items. Any neural activity tied to the presentation of individual items and predictive of encoding success would have survived our shuffling procedure and thus should have resulted in a list-level SME even for shuffled lists. Instead these findings suggest that relevant endogenous factors vary at the time scale of multiple item presentations. Averaging brain activity across encoding periods within a list thus yields a signal that is strongly predictive of list-level recall performance, because items that are studied together are studied in similar “cognitive states.” These findings raise the questions about the nature of the relevant endogenous factors producing these states. The prominent negative correlation between recall performance and α power (shown in Figures 1c, 3b, and 4b) could suggest that the endogenous factors that drive the SMEs reflect attentional engagement during memory encoding (Sadaghiani & Kleinschmidt, 2016). According to this interpretation, SMEs would not specifically index mnemonic encoding processes and should generalize to other tasks without memory tests. Further work is required to establish the extent to which SMEs reflect general attentional processes or specifically relate to successful memory encoding. Within the multivariate approach introduced here, this question could be addressed by contrasting decoding and cross-decoding performance of multivariate models applied to different tasks (Weidemann et al., 2019).

Because SMEs have been demonstrated in tasks other than free recall, and for various measures of brain activity (Hanslmayr & Staudigl, 2014; Fernandez, Brewer, Zhao, Glover, & Gabrieli, 1999; Otten, Henson, & Rugg, 2002; Schott et al., 2011), future work will need to address the question of how endogenous neural variation underlies memory encoding outside of our experimental setting. The fact that substantial SMEs remained after accounting for a comprehensive set of external variables may appear in conflict with findings that encoding task manipulations can affect the specific form of SMEs, at least for recognition memory (Kamp, Bader, & Mecklinger, 2017; Summerfield & Mangels, 2006; Otten & Rugg, 2001; Staudigl & Hanslmayr, 2013; Fellner et al., 2013). Here we show that in the absence of direct manipulations of how study items are presented or processed, SMEs mainly reflect endogenous factors with relatively modest contributions from external factors, at least for EEG activity in a free recall task.

Our findings align with reports of sequential dependencies in human performance (Kahana et al., 2018; Gilden et al., 1995; Mueller & Weidemann, 2008; Verplanck et al., 1952) as well as with those of slow endogenous neural fluctuations that drive variability in evoked brain activity and overt behavior (Monto et al., 2008; Schroeder & Lakatos, 2009; Arieli et al., 1996; Fox et al., 2005, 2007; Fox & Raichle, 2007; Raichle, 2015). Previous investigations of endogenous variability in neural activity and performance have relied on exact repetitions of stimuli across many experimental trials to limit variability in external factors. To study the effects of endogenous variability on recall performance, we took a complementary approach by statistically removing the effects of a comprehensive set of external factors. Despite the differences in methodologies and tasks, the conclusions are remarkably consistent in establishing an important role for slowly varying fluctuations in neural activity as drivers of variability in human cognition.

Because encoding and retrieval processes jointly determine mnemonic success, it is notoriously difficult to study either process in isolation. The assessment of encoding-related brain activity as a function of subsequent memory performance offers a powerful tool for isolating neural processes specifically underlying memory formation. As typically used, however, this method conflates external factors that predict subsequent memory (e.g., item complexity) and endogenously varying neural processes. Here we used two new methods to deconfound these factors: First, we used a statistical model to control for external factors and examined the SME on residual performance measures. Second, we introduced a new list-level SME and a session-level resampling control procedure that identifies encoding-related neural activity that varies at the time-scale of entire list presentations. Both approaches showed that endogenous neural activity dominates the subsequent memory effect, highlighting its effectiveness for the study of cognitive processes associated with memory acquisition.

## Supporting information

Supplementary table of stimuli

## References

Arieli, A., Sterkin, A., Grinvald, A., & Aertsen, A. (1996). Dynamics of ongoing activity: Explanation of the large variability in evoked cortical responses. Science, 273, 1868–1871. doi: 10.1126/science.273.5283.1868

Cohen, J. (1988). Statistical power analysis for the behavioral sciences. Lawrence Erlbaum Associates.

DeLosh, E. L., & McDaniel, M. A. (1996). The role of order information in free recall: Application to the word-frequency effect. Journal of Experimental Psychology: Learning, Memory, and Cognition, 22, 1136–1146. doi: 10.1037/0278-7393.22.5.1136

Ezzyat, Y., Wanda, P. A., Levy, D. F., Kadel, A., Aka, A., Pedisich, I., … Kahana, M. J. (2018). Closed-loop stimulation of temporal cortex rescues functional networks and improves memory. Nature Communications, 9(1). doi: 10.1038/s41467-017-02753-0

Fellner, M.-C., Bäuml, K.-H. T., & Hanslmayr, S. (2013). Brain oscillatory subsequent memory effects differ in power and longrange synchronization between semantic and survival processing. NeuroImage, 79, 361–370. doi: 10.1016/j.neuroimage.2013.04.121

Fernandez, G., Brewer, J. B., Zhao, Z., Glover, G. H., & Gabrieli, J. D. (1999). Level of sustained entorhinal activity at study correlates with subsequent cued-recall preformance: a functional magnetic resonance imaging study with high acquisition rate. Hippocampus, 9, 35–44. doi: 10.1002/(SICI)1098-1063(1999)9:1<35::AID-HIPO4>3.0.CO;2-Z

Fox, M. D., & Raichle, M. E. (2007). Spontaneous fluctuations in brain activity observed with functional magnetic resonance imaging. Nature Reviews Neuroscience, 8, 700–711. doi: 10.1038/nrn2201

Fox, M. D., Snyder, A. Z., Vincent, J. L., & Raichle, M. E. (2007). Intrinsic fluctuations within cortical systems account for intertrial variability in human behavior. Neuron, 56, 171–184. doi: 10.1016/j.neuron.2007.08.023

Fox, M. D., Snyder, A. Z., Zacks, J. M., & Raichle, M. E. (2005). Coherent spontaneous activity accounts for trial-to-trial variability in human evoked brain responses. Nature Neuroscience, 9, 23–25. doi: 10.1038/nn1616

Gilden, D., Thornton, T., & Mallon, M. (1995). 1/f noise in human cognition. Science, 267, 1837–1839. doi: 10.1126/science.7892611

Gramfort, A., Luessi, M., Larson, E., Engemann, D. A., Strohmeier, D., Brodbeck, C., … Hämäläinen, M. (2013). MEG and EEG data analysis with MNE-python. Frontiers in Neuroscience, 7, 267. doi: 10.3389/fnins.2013.00267

Gramfort, A., Luessi, M., Larson, E., Engemann, D. A., Strohmeier, D., Brodbeck, C., … Hämäläinen, M. S. (2014). MNE software for processing MEG and EEG data. NeuroImage, 86, 446–460. doi: 10.1016/j.neuroimage.2013.10.027

Guderian, S., Schott, B. H., Richardson-Klavehn, A., & Duzel, E. (2009). Medial temporal theta state before an event predicts episodic encoding success in humans. Proceedings of the National Academy of Sciences, 106(13), 5365–5370. doi: 10.1073/pnas.0900289106

Hanslmayr, S., Spitzer, B., & Bäuml, K.-H. (2009). Brain oscillations dissociate between semantic and nonsemantic encoding of episodic memories. Cerebral Cortex, 19, 1631–1640. doi: 10.1093/cercor/bhn197

Hanslmayr, S., & Staudigl, T. (2014). How brain oscillations form memories — a processing based perspective on oscillatory subsequent memory effects. NeuroImage, 85, 648–655. doi: 10.1016/j.neuroimage.2013.05.121

Healey, M. K., Crutchley, P., & Kahana, M. J. (2014). Individual differences in memory search and their relation to intelligence. Journal of Experimental Psychology: General, 143, 1553–1569. doi: 10.1037/a0036306

Healey, M. K., & Kahana, M. J. (2014). Is memory search governed by universal principles or idiosyncratic strategies? Journal of Experimental Psychology: General, 143, 575–596. doi: 10.1037/a0033715

Healey, M. K., & Kahana, M. J. (2018). Age-related changes in the dynamics of memory encoding processes provide a biomarker of successful aging. Manuscript Submitted for publication.

Kahana, M. J., Aggarwal, E. V., & Phan, T. D. (2018). The variability puzzle in human memory. Journal of Experimental Psychology: Learning, Memory, and Cognition. doi: 10.1037/xlm0000553

Kamp, S.-M., Bader, R., & Mecklinger, A. (2017). ERP subsequent memory effects differ between inter-item and unitization encoding tasks. Frontiers in Human Neuroscience, 11. doi: 10.3389/fnhum.2017.00030

Kim, H. (2011). Neural activity that predicts subsequent memory and forgetting: A meta-analysis of 74 fMRI studies. NeuroImage, 54(3), 2446–2461. doi: 10.1016/j.neuroimage.2010.09.045

Lohnas, L. J., & Kahana, M. J. (2013). Parametric effects of word frequency in memory for mixed frequency lists. Journal of Experimental Psychology: Learning, Memory, and Cognition, 39, 1943–1946. doi: 10.1037/a0033669

Lohnas, L. J., Polyn, S. M., & Kahana, M. J. (2015). Expanding the scope of memory search: Modeling intralist and interlist effects in free recall. Psychological Review, 122, 337–363. doi: 10.1037/a0039036

Merritt, P. S., DeLosh, E. L., & McDaniel, M. A. (2006). Effects of word frequency on individual-item and serial order retention: Tests of the order-encoding view. Memory & Cognition, 34, 1615–1627. doi: 10.3758/bf03195924

Monto, S., Palva, S., Voipio, J., & Palva, J. M. (2008). Very slow EEG fluctuations predict the dynamics of stimulus detection and oscillation amplitudes in humans. Journal of Neuroscience, 28, 8268–8272. doi: 10.1523/jneurosci.1910-08.2008

Mueller, S. T., & Weidemann, C. T. (2008). Decision noise: An explanation for observed violations of signal detection theory. Psychonomic Bulletin & Review, 15, 465–494. doi: 10.3758/pbr.15.3.465

Murdock, B. B., Jr. (1962). The serial position effect of free recall. Journal of Experimental Psychology, 64, 482–488. doi: 10.1037/h0045106

Otten, L. J., Henson, R. N. A., & Rugg, M. D. (2002, oct). State-related and item-related neural correlates of successful memory encoding. Nature Neuroscience, 5(12), 1339–1344. doi: 10.1038/nn967

Otten, L. J., Quayle, A. H., Akram, S., Ditewig, T. A., & Rugg, M. D. (2006). Brain activity before an event predicts later recollection. Nature Neuroscience, 9(4), 489–491. doi: 10.1038/nn1663

Otten, L. J., & Rugg, M. D. (2001). Electrophysiological correlates of memory encoding are task-dependent. Cognitive Brain Research, 12, 11–18. doi: 10.1016/s0926-6410(01)00015-5

Paller, K. A., & Wagner, A. D. (2002). Observing the transformation of experience into memory. Trends in Cognitive Sciences, 6, 93–102. doi: 10.1016/s1364-6613(00)01845-3

Park, H., & Rugg, M. D. (2010). Prestimulus hippocampal activity predicts later recollection. Hippocampus, 20, 24–28. doi: 10.1002/hipo.20663

Pedregosa, F., Varoquaux, G., Gramfort, A., Michel, V., Thirion, B., Grisel, O., … Duchesnay, E. (2011). Scikit-learn: Machine learning in Python. Journal of Machine Learning Research, 12, 2825–2830.

Raichle, M. E. (2015). The restless brain: how intrinsic activity organizes brain function. Philosophical Transactions of the Royal Society B: Biological Sciences, 370, 20140172. doi: 10.1098/rstb.2014.0172

Sadaghiani, S., & Kleinschmidt, A. (2016). Brain networks and α-oscillations: Structural and functional foundations of cognitive control. Trends in Cognitive Sciences, 20, 805–817. doi: 10.1016/j.tics.2016.09.004

Schott, B. H., Wüstenberg, T., Wimber, M., Fenker, D. B., Zierhut, K. C., Seidenbecher, C. I., … Richardson-Klavehn, A. (2011). The relationship between level of processing and hippocampalcortical functional connectivity during episodic memory formation in humans. Human Brain Mapping, 34, 407–424. doi: 10.1002/hbm.21435

Schroeder, C. E., & Lakatos, P. (2009). Low-frequency neuronal oscillations as instruments of sensory selection. Trends in Neurosciences, 32, 9–18. doi: 10.1016/j.tins.2008.09.012

Siegel, L. L., & Kahana, M. J. (2014). A retrieved context account of spacing and repetition effects in free recall. Journal of Experimental Psychology: Learning, Memory, and Cognition, 40, 755–764. doi: 10.1037/a0035585

Staudigl, T., & Hanslmayr, S. (2013). Theta oscillations at encoding mediate the context-dependent nature of human episodic memory. Current Biology, 23(12), 1101–1106. doi: 10.1016/j.cub.2013.04.074

Summerfield, C., & Mangels, J. A. (2006). Dissociable neural mechanisms for encoding predictable and unpredictable events. Journal of Cognitive Neuroscience, 18, 1120–1132. doi: 10.1162/jocn.2006.18.7.1120

Sweeney-Reed, C. M., Zaehle, T., Voges, J., Schmitt, F. C., Buentjen, L., Kopitzki, K., … Rugg, M. D. (2016). Pre-stimulus thalamic theta power predicts human memory formation. NeuroImage, 138, 100–108. doi: 10.1016/j.neuroimage.2016.05.042

Urgolites, Z. J., Wixted, J. T., Goldinger, S. D., Papesh, M. H., Treiman, D. M., Squire, L. R., & Steinmetz, P. N. (2020). Spiking activity in the human hippocampus prior to encoding predicts subsequent memory. Proceedings of the National Academy of Sciences, 117(24), 13767–13770. doi: 10.1073/pnas.2001338117

Verplanck, W. S., Collier, G. H., & Cotton, J. W. (1952). Nonindependence of successive responses in measurements of the visual threshold. Journal of Experimental Psychology, 44, 273–282. doi: 10.1037/h0054948

Warriner, A. B., Kuperman, V., & Brysbaert, M. (2013). Norms of valence, arousal, and dominance for 13,915 English lemmas. Behavior research methods, 45(4), 1191–1207.

Weidemann, C. T., & Kahana, M. J. (2016). Assessing recognition memory using confidence ratings and response times. Royal Society Open Science, 3, 150670. doi: 10.1098/rsos.150670

Weidemann, C. T., & Kahana, M. J. (2019). Dynamics of brain activity reveal a unitary recognition signal. Journal of Experimental Psychology: Learning, Memory, and Cognition, 45, 440–451. doi: 10.1037/xlm0000593

Weidemann, C. T., Kragel, J. E., Lega, B. C., Worrell, G. A., Sperling, M. R., Sharan, A. D., … Kahana, M. J. (2019). Neural activity reveals interactions between episodic and semantic memory systems during retrieval. Journal of Experimental Psychology: General, 148, 1–12. doi: 10.1037/xge0000480

Weidemann, C. T., Mollison, M. V., & Kahana, M. J. (2009). Electrophysiological correlates of high-level perception during spatial navigation. Psychonomic Bulletin & Review, 16, 313–319. doi: 10.3758/pbr.16.2.313

