## Supplementary table of stimuli for "Neural measures of subsequent memory reflect endogenous variability in cognitive function"

### Word pool

ACTOR
ACTRESS
AGENT
AIRPLANE
AIRPORT
ANKLE
ANTLER
APPLE
APRON
ARM
ARMY
ASIA
ATLAS
ATOM
AUTHOR
AWARD
BABY
BACKBONE
BACON
BADGE
BALLOON
BANJO
BANK
BANKER
BANQUET
BARLEY
BARREL
BASEMENT
BATHTUB
BEAKER
BEAST
BEAVER
BEEF
BELLY
BIKE
BINDER
BISON
BLACKBOARD
BLADE
BLENDER
BLOCKADE
BLOUSE
BLUEPRINT
BOARD
BODY
BOUQUET
BOX
BOYFRIEND
BRACES
BRAKE
BRANCH
CLOCK
CLOTHES
CLOUD
COBRA
COCKTAIL
COCOON
COD
COFFEE
COIN
COLLEGE
COLONEL
COMET
COMPASS
CONCERT
CONTRACT
CONVICT
COOK
COOKBOOK
CORAL
COSTUME
COTTAGE
COUCH
COUNTRY
COUNTY
COURSE
COUSIN
COWBOY
CRAB
CRATER
CRAYON
CREATURE
CREVICE
CRIB
CRICKET
CRITIC
CROSS
CROWN
CRUTCH
CUPBOARD
CURTAIN
CUSTARD
CYCLONE
DAISY
DANCER
DANDRUFF
DASHBOARD
DAUGHTER
DENIM
DENTIST
DIME
DINER
FRANCE
FRECKLE
FREEZER
FRIAR
FRIEND
FRUIT
FUNGUS
GALLON
GANGSTER
GARBAGE
GARDEN
GARLIC
GAVEL
GAZELLE
GHETTO
GIFT
GIRL
GLASS
GLOBE
GLOVE
GOBLIN
GRAPE
GRAVE
GRAVEL
GREASE
GRILL
GRIZZLY
GROUND
GUARD
GUITAR
GYMNAST
HAMPER
HAND
HANDBAG
HARP
HATCHET
HAWK
HEADBAND
HEART
HEDGE
HELMET
HERO
HIGHWAY
HIKER
HONEY
HOOD
HOOK
HORNET
HORSE
HOSTESS
HOUND
LIZARD
LODGE
LOFT
LONDON
LOVER
LUGGAGE
LUMBER
LUNCH
MACHINE
MAILBOX
MAILMAN
MAMMAL
MAPLE
MARINE
MARKER
MARKET
MARROW
MARS
MARSH
MASK
MATCH
MATTRESS
MEAT
MEDAL
MESSAGE
MILDEW
MILK
MISSILE
MISTER
MONEY
MONSTER
MOP
MOTEL
MOTOR
MUFFIN
MUMMY
MUSTARD
NAPKIN
NECKLACE
NEUTRON
NIGHTGOWN
NOMAD
NOTEBOOK
NOVEL
NURSE
OFFICE
OINTMENT
OMELET
ONION
ORANGE
ORCHID
PLAYGROUND
PLAZA
PLIERS
PLUTO
POCKET
POET
POISON
POLICE
POPCORN
PORK
PORTRAIT
POSSUM
POSTAGE
POWDER
PREACHER
PRIMATE
PRINCE
PRINCESS
PROTON
PUDDING
PUDDLE
PUPIL
PUPPY
QUAIL
QUARTER
QUEEN
RABBIT
RACKET
RADISH
RAFT
RATTLE
RAZOR
REBEL
RECEIPT
RECORD
RELISH
REPORT
RIFLE
RIVER
ROBBER
ROBIN
ROBOT
ROCKET
ROD
ROOSTER
RUG
RUST
SADDLE
SALAD
SALMON
SALT
SPIDER
SPONGE
SPOOL
SPOON
SPOUSE
STAKE
STALLION
STAMP
STAPLE
STAR
STATUE
STICKER
STOMACH
STONE
STOVE
STREAM
STUDENT
SUBWAY
SUITCASE
SUMMIT
SUNRISE
SUNSET
SUPPER
SURVEY
SUSPECT
SWAMP
SWIMMER
SWITCH
SWORD
TABLE
TABLET
TART
TAXI
TEACHER
TEMPLE
TERMITE
THIEF
THREAD
THRONE
TILE
TOASTER
TOMBSTONE
TORTOISE
TOURIST
TRACTOR
TRANSPLANT
TREAT
TRENCH
TRIBE
TROMBONE
TROUT

BRANDY
BREAST
BRICK
BRIEFCASE
BROOK
BROTHER
BUBBLE
BUCKET
BUG
BUGGY
BULLET
BUNNY
BUREAU
BURGLAR
BUTCHER
CABBAGE
CABIN
CAFE
CAMEL
CANAL
CANDY
CANYON
CAPTIVE
CARRIAGE
CARROT
CASHEW
CASHIER
CASKET
CATCHER
CATTLE
CEILING
CELLAR
CHAMPAGNE
CHAPEL
CHAUFFEUR
CHEMIST
CHEST
CHILD
CHIPMUNK
CHURCH
CIGAR
CITRUS
CLAM
CLAMP
CLIMBER
DIVER
DOLPHIN
DONKEY
DONOR
DORM
DOUGHNUT
DRAGON
DRAWING
DRESS
DRESSER
DRILL
DRINK
DRIVER
DRUG
DUST
DUSTPAN
EAGLE
EGYPT
ELBOW
EMPIRE
EUROPE
EXPERT
EYELASH
FARMER
FEMALE
FIDDLE
FILM
FINGER
FIREMAN
FIREPLACE
FLAG
FLASHLIGHT
FLASK
FLEET
FLESH
FLIPPER
FLOWER
FLUTE
FOOT
FOOTBALL
FOREHEAD
FOREST
FOX
FRAGRANCE
FRAME
HUMAN
HUSBAND
ICEBERG
ICING
IDOL
IGLOO
INFANT
INMATE
ISLAND
ITEM
JAPAN
JEANS
JELLO
JELLY
JOURNAL
JUDGE
JUGGLER
JUNGLE
JURY
KEEPER
KETCHUP
KIDNEY
KITCHEN
KLEENEX
KNAPSACK
KNIFE
LABEL
LACE
LADDER
LADY
LAGOON
LAKE
LAMP
LAPEL
LASER
LAVA
LEADER
LEG
LEOPARD
LETTUCE
LIGHTNING
LILY
LION
LIPSTICK
LIVER
OUTDOORS
OUTFIT
OUTLAW
OX
OYSTER
OZONE
PACKAGE
PADDING
PADDLE
PAIL
PALACE
PANTHER
PAPER
PARENT
PARROT
PARSLEY
PARTNER
PASSAGE
PASTA
PASTRY
PATIENT
PATROL
PEACH
PEANUT
PEBBLE
PECAN
PEDAL
PENGUIN
PEPPER
PERCH
PERFUME
PERMIT
PIANO
PICNIC
PICTURE
PIGEON
PIGMENT
PILOT
PIMPLE
PISTOL
PISTON
PIZZA
PLAID
PLASTER
PLATE
SANDWICH
SAUSAGE
SCALLOP
SCALPEL
SCARECROW
SCARF
SCISSORS
SCOTCH
SCRIBBLE
SCULPTURE
SEAFOOD
SEAGULL
SEAL
SERVANT
SERVER
SHARK
SHELF
SHELTER
SHERIFF
SHIRT
SHORTCAKE
SHORTS
SHOULDER
SHOVEL
SHRUB
SIBLING
SIDEWALK
SILK
SISTER
SKETCH
SKILLET
SKIRT
SLIDE
SLIME
SLOPE
SLUG
SMOG
SNACK
SNAIL
SNAKE
SODA
SOFTBALL
SPACE
SPARROW
SPHINX
TRUCK
TUBA
TUNNEL
TURKEY
TURNIP
TURTLE
TUTU
TWEEZERS
TWIG
TWISTER
TYPIST
ULCER
UMPIRE
UNCLE
VAGRANT
VALLEY
VALVE
VELVET
VENUS
VICTIM
VIKING
VIRUS
WAGON
WAITER
WAITRESS
WARDROBE
WASHER
WASP
WHISKERS
WHISTLE
WIDOW
WIFE
WINDOW
WITNESS
WOMAN
WORKER
WORLD
WRENCH
WRIST
XEROX
YACHT
YARN
YOLK
ZEBRA
ZIPPER
